# Reduced glycolysis links resting zone chondrocyte proliferation in the growth plate

**DOI:** 10.1101/2023.01.18.524550

**Authors:** Tatsuya Kobayashi, Cameron Young, Wen Zhou, Eugene P. Rhee

## Abstract

A gain-of-function mutation of the chondrocyte-specific microRNA, miR-140-5p, encoded by the MIR140 gene, causes spondyloepiphyseal dysplasia, Nishimura type (SEDN, also known as SED, MIR140 type; MIM, 611894). We reported that a mouse model for SEDN showed a unique growth plate phenotype that is characterized by an expansion of the resting zone of the growth plate and an increase in resting chondrocytes, of which the mechanism of regulation is poorly understood. We found that the miR-140 mutant chondrocytes showed a significant reduction of Hif1a, the master transcription factor that regulates energy metabolism in response to hypoxia. Based on this finding, we hypothesized that energy metabolism plays a regulatory role in resting chondrocyte proliferation and growth plate development. In this study, we show that suppression of glycolysis via LDH ablation causes an expansion of the resting zone and skeletal developmental defects. We have also found that reduced glycolysis results in reduced histone acetylation in the miR-140 mutant as well as LDH-deficient chondrocytes likely due to the reduction in acetyl-CoA generated from mitochondria-derived citrate. Reduction in acetyl-CoA conversion from citrate by deleting Acly caused an expansion of the resting zone and a similar gross phenotype to LDH-deficient bones without inducing energy deficiency, suggesting that the reduced acetyl-CoA, but not the ATP synthesis deficit, is responsible for the increase in resting zone chondrocytes. Comparison of the transcriptome between LDH-deficient and Acly-deficient chondrocytes also showed overlapping changes including upregulation in Fgfr3. We also confirmed that overexpression of an activation mutation of Ffgr3 causes an expansion of resting zone chondrocytes. These data demonstrate the association between reduced glycolysis and an expansion of the resting zone and suggest that it is caused by acetyl-CoA deficiency, but not energy deficiency, possibly through epigenetic upregulation of FGFR3 signaling.

## Introduction

Skeletal development requires tightly coordinated proliferation and differentiation of growth plate chondrocytes, and aberrations of this process lead to diverse types of skeletal dysplasia ^1^. In the growth plate, chondrocytes in the resting zone slowly proliferate, differentiate into vigorously proliferating columnar chondrocytes, then further into hypertrophic chondrocytes which drive longitudinal bone growth ^2^. The resting zone chondrocytes are the most immature chondrocytes in the growth plate and are considered to serve as tissue-specific stem/progenitor cells; however, limited knowledge is available regarding the mechanisms by which the proliferation and differentiation of resting chondrocytes are regulated.

We previously reported that a mouse model for spondyloepiphyseal dysplasia, Nishimura type (SEDN, also known as SED, MIR140 type; MIM, 611894), caused by a gain-of-function mutation of microRNA-140-5p (miR-140-5p) that is encoded by the MIR140 gene, show a unique growth plate phenotype where the growth plate is expanded mainly due to the increase in the number of resting zone chondrocytes ^3^. The RNA-seq data suggested reduced hypoxic adaptation in the mutant chondrocytes, including the reduced expression of the master transcription factor, hypoxia-inducible factor alpha (Hif1a). Since a binding site of mutant miR-140 was found in the coding sequence of *Hif1a*, it was suggested that the mutant miR-140-5p directly suppresses *Hif1a* expression. Hif1a is an essential transcription factor for cell survival and proliferation of growth plate chondrocytes ^4; 5^. In hypoxic environments, like in developing growth plates, Hif1a protein is stabilized, and it stimulates glycolysis and suppresses mitochondrial metabolism by inhibiting pyruvate entry to mitochondria^6^. Live cells actively metabolize diverse molecules to meet their demands for energy and biomolecule synthesis. At the same time, various metabolites produced during energy metabolism are known to have regulatory roles in diverse cell functions ^7; 8^. Therefore, in addition to energy deficiency, it is also possible that the altered metabolic state of the mutant miR-140-5p-expressing chondrocytes causes the deregulation of signaling pathways and gene expression critical for controlling chondrocyte proliferation and differentiation.

Adenosine triphosphate (ATP), cells’ energy currency, is generated primarily by glucose catabolism. During glycolysis, a single molecule of glucose is broken down into pyruvate, while generating 2 molecules of ATPs. Pyruvate is further transported into mitochondria and metabolized into acetyl CoA in the tricarboxylic acid (TCA) cycle to generate electron donors. The electron transport chain then creates proton gradients, which drive ATP synthase to efficiently generate ATPs while consuming oxygen, through the process called oxidative phosphorylation (OXPHOS). Pyruvate can be directly metabolized in the cytoplasm into lactic acid by the enzyme, lactate dehydrogenase (LDH), which is encoded by the Ldha, Ldhb, Ldhc, and Ldhd genes in mice. This process is considered to be necessary to produce nicotinamide adenine dinucleotide (NAD^+^), which is an essential co-substrate for glyceraldehyde-3-phosphate dehydrogenase (Gapdh), a key enzyme for glycolysis ^9^. Therefore, by reducing LDH activity, glycolysis can be suppressed at an upstream step of the glycolytic cascade.

The growth plate is an avascular tissue, and therefore, it is mostly under a hypoxic condition, particularly during development. With the limited availability of oxygen, chondrocytes primarily rely on glycolysis for energy production^10^. This notion is supported by the fact that inhibition of glucose uptake in chondrocytes resulted in reduced chondrocyte proliferation and developmental defects of endochondral bones ^11^. Although glycolysis is the major mechanism for ATP production in chondrocytes, mitochondrial metabolism still plays an important role in growth plate chondrocytes, as conditional deletion of Tfam, a transcription factor critical for mitochondrial genome transcription, impairs normal endochondral bone development ^12^. In addition to energy production, mitochondria play important regulatory roles by providing various bioactive molecules. For example, citrate in the TCA cycle is exported to the cytoplasm and converted back into acetyl-CoA, which is the primary source for lipid synthesis as well as the donor of the acetyl group for protein modification^7; 8^. It is known that Ac-CoA availability directly determines the acetylation level of proteins, including histones, the most abundantly acetylated group of proteins in the cell ^13^.

Given the importance of glucose metabolism in chondrocytes and the unique growth plate phenotype of the SEDN mouse model that shows reduced Hif1a expression and altered metabolic gene expression, we hypothesize that energy metabolism has regulatory roles in growth plate development.

In this study, we deleted the major LDH genes, *Ldha* and *ldhb* to suppress glycolysis in chondrocytes. We show that LDH ablation causes skeletal developmental defects with increased resting chondrocyte proliferation. LDH deficiency causes compensatory upregulation of OXPHOS, leading to reductions of cytoplasmic citrate, Ac-CoA, and acetylated histone levels. We also show that reduction in Ac-CoA by *Acly* deletion leads to a growth plate and molecular phenotype similar to that of LDH deficiency without inducing energy deficit, suggesting that the reduced level of Ac-CoA, but not ATP synthesis, is responsible for resting chondrocyte proliferation.

## Material and methods

### Mice

The animal experiments were approved by the Institutional Animal Care and Use Committee (IACUC) at Massachusetts General Hospital.

Floxed *Ldha* mice were previously described ^14^ and purchased from the Jackson Laboratory. Floxed Acly mice were previously described ^15^. Transgenic mice with a construct of Cre-dependent expression of a constitutively active Fgfr3 mutant (Tg(CAG-Fgfr3*K650E,-EGFP)10Jheb) ^16^ were purchased from the Jackson Laboratory. *Col2-Cre* transgenic mice and Sox9-CreER transgenic mice were described ^17; 18^. *Ldhb* knockout (*Ldhb*^*-/-*^) mice were generated by the CRISPR-mediated *in vivo* genome editing, i-GONAD^19^ using the CD-1 outbred strain (Charles River Laboratory). Briefly, at 0.7 day post coitum, the oviduct of female mice was exposed through a dorsal incision. Approximately 1 ul of CRISPR solution containing 1 ug/ul of Cas9 (Sigma or IDT) premixed with 10 mM two-piece guide RNA (crRNA and tracrRNA, gRNA) (IDT), was injected into the oviduct either by puncturing the oviductal wall or through the infundibulum. Then, *in vivo* electroporation was performed using the square pulse generator, BTX-820 with the electrode tweezers, CUY652P2.5×4 (Nepa Gene, Japan) with the setting of 50V, 5msec each pulse, 8 pulses, with 0.5 cm electrode gap. Pups were genotyped for the desired modification, then mated with wildtype mice to establish a line. The Ldhb gene was deleted by cleaving the genome at two sites using two gRNAs mixed with Cas9 to remove a 15-kb long genomic region containing exons encoding more than 90% of the *Ldhb* coding sequence including the entire active site. The spacer sequences of these crRNAs are Ldhb-L 5’-CGCAAUGAGCUUCUCCUUAA-3’ and Ldhb-R 5’-ACGAUGAGGUCGCUCAGCUC-3’. Genotyping of the Ldhb gene was performed by PCR using the wildtype-specific primer set, Ldhb-L 5’-CCTTCAGGGCTTCTGTTGAG -3’ and Ldhb-R 5’-CATGTCAGGGAAGAAGCAAA -3’, and the KO-specific primer set, Ldhb-L and Ldhb-Wt 5’-CGCCCACTACAGTGATCTTG -3’. Mir140UGCG mice that express a gain-of-function mutation of the miR-140-5p gene (Mir140G, miR-140-5pG) ^3^ with additional mutations were described ^20^. The additional mutations (the first nucleotide and last two nucleotides of miR-140-5p sequence of *Mir140*) facilitate the loading of miR-140-5p onto Argonaute proteins, and thereby enhances the expression of miR-140-5p-G and reduces miR-140-3p expression. These mutations do not alter the seed sequence and therefore do not significantly change its target RNA repertoire. Mir140UGCG mice show a greater gain-of-function effect of miR-140 mutation than Mir140G at the heterozygous state.

We found that *Col2-Cre:Ldha*^*fl/fl*^ mice in the C57/B6 background were mostly perinatally lethal but that they survived postnatally in a mixed background of CD1 and C57/B6. Therefore, the experiments in this study were performed in a CD1-dominant congenic background.

### Histological analyses

Mice were sacrificed at indicated ages. Tissues were dissected and fixed in 10% Formaldehyde/PBS for histological analysis. Decalcification was performed in EDTA for 2 weeks on a necessary basis, and then tissues were processed in paraffin. Sections were cut, stained with Hematoxylin and Eosin (H & E), or subjected to other analyses according to the standard protocols.

Immunofluorescence was performed according to the protocol described in the data sheets of the anti-acetylated histone 3 lysine 27 (H3K27Ac) antibody (# 8173, Cell Signaling Technology) and anti-acetylate histone 3 lysine 9 (H3K9Ac) antibody (#9649, Cell Signaling Technology).

The fluorescent labeled secondary antibody (Alexa 568 anti-rabbit goat antibody, A-11011) was purchased from ThermoFisher Scientific.

EdU (5-ethynyl-2′-deoxyuridine) labeling assay was performed according to the instruction of EdU Assay / EdU Staining Proliferation Kit (ab219801, Abcam). Proliferating cells were labeled by injecting 10mg/kg BW EdU to mice intraperitoneally 2 hours before sacrifice. Tissues were fixed, paraffin-processed, and subjected to EdU staining and DAPI nuclear staining.

### Primary cell Isolation and cell culture

Mice were sacrificed at P10. Primary rib chondrocytes were isolated from these mice as previously described with some modifications ^21^. After overnight collagenase digestion, the cells were placed in 2 mL tubes, centrifuged, and the collagenase was removed from the cells. Cells were resuspended in DMEM, 10% FBS, and 1% P/S. Cells were passed through a 0.40 um strainer (Corning) into a 6-well plate (Corning). Cells were cultured in DMEM-containing 10% FBS and antibiotics, counted, and replated for subsequent assays. Primary growth plate chondrocytes were isolated as previously reported with modifications ^21^. Cells were cultured in DMEM containing 4.5g/L of glucose and 10% fetal bovine serum (FBS) without lactate in a humidified 37ºC incubator, otherwise specified.

### RNA-seq analysis

RNA-seq was performed using polyA-enriched RNA from primary rib chondrocytes isolated from P5 mice of *Col2-Cre:Ldha*^*fl/fl*^*:Ldhb*^*+/-*^ and wildtype littermate control or from E18.5 rib chondrocytes of *Col2-Cre:Acly* ^*fl/fl*^ embryos and *Cre*-negative littermate control. RNA-seq was performed by Beijing Genome Institute (BGI). RNA-seq data is available at Gene Expression Omnibus (GEO) with the accession number GSE192971.

### Immunoblot analysis

Immunoblot was performed according to a standard procedure using 4%-20% gradient SDS polyacrylamide gels (Genscript). Cells were directly lysed in 4X Laemmli gel loading buffer (Boston BioProducts), and protein was transferred onto nitrocellulose membranes using the Genie Blotter (Idea Scientific). The anti-Actb (#8457), anti-H3K27Ac (# 8173), and anti-H3K9Ac (#9649) antibodies were purchased from Cell Signaling Technology.

### Seahorse analysis

Net activities of glycolysis and mitochondrial respiration were assessed using the mitostress test using the Seahorse XFe96 analyzer (Agilent) according to the manufacturer’s instruction. Briefly, primary chondrocytes or bone marrow stromal cells, prepared as described above were plated at the near confluence and cultured overnight in a growth medium (DMEM containing 10% FBS) before subjecting them to analysis.

### Lipid analysis

Metabolites were assayed as previously described ^22; 23^. In brief, Lipids were extracted from primary chondrocytes in monolayer culture in isopropanol and quantified using C8 chromatography and nontargeted, positive ion mode MS analysis on an Exactive Plus Orbitrap Mass Spectrometer. Identification of known metabolites was achieved by matching retention times and mass/charge ratio (*m/z*) to synthetic mixtures of reference compounds and characterized pooled plasma reference samples. Values were normalized by the cell mass. Lipid species were categorized into 6 groups. The average value of each lipid species was calculated, and the fold difference between control and mutant chondrocytes was plotted.

### Statistical Analysis

Statistical analysis was performed using Prism 9 (GraphPad)

## Results

### Enhanced Expression of a Gain-of-Function Mutation of Mir140 Causes expansion of the Resting Zone of the Growth Plate

The Mir140 gene encodes three microRNAs (miRNAs), miR-140-5p, miR-140-3p.1 and miR-140-3p.2 that are generated from a single pre-miR-140 hairpin precursor upon cleavage by the RNase III, Dicer. Then either the -5p or a -3p miRNA is selected and loaded onto Argonaute (Ago) proteins ^24^. In chondrocytes, the abundance of miR-140-3p miRNAs, miR-140-3p.1 and - 3p.2, is about 10 times greater than that of miR-140-5p likely due to asymmetric selection of miR-140 duplex strands during Ago loading. This strand selection appears to generally follow simple rules; the strand that starts uridine (U) or adenosine (A) is preferred for loading that those with guanosine (G) or cytidine (C), and the strand of which 5’ end is located at the more thermodynamically unstable side of a duplex is preferred ^25; 26^.

We previously reported that a single nucleotide substitution of the MIR140 in humans caused ultra-rare skeletal dysplasia, spondyloepiphyseal dysplasia, Nishimura type (SEDN, also known as SED, MIR140 type; MIM, 611894)^3^. In this condition, the single nucleotide substitution at the second nucleotide of miR-140-5p miRNA, from adenosine (A) to guanosine (G), causes a gain-of-function (GOF) effect, leading to skeletal dysplasia both in humans and mice. In the mouse model carrying the identical substitution as in patients, the resting zone of the growth plate was expanded, causing a delay in ossification and development of the epiphysis.

The *Mir140* GOF mutation showed a dose-dependent effect; homozygotes show a more severe phenotype than heterozygotes. To create a more robust mouse model, we introduced minor changes in the nucleotides at the 5’ end of miR-140-5p and -3p to increase the expression level of miR-140-5p; the substitutions from C to U at the 5’-end of miR-140-5p and the change from UA to CG at the 5’-end of miR-140-3p increase the Ago loading efficiency of the -5p strand, thus enhances the GOF effect ^20^ (Figure 1A). As expected, the mutant mice expressing the GOF- mutant miR-140-5p (miR-140-5pG) with enhanced expression (*Mir140UGCG*) showed a significant expansion of the growth plate even in the heterozygous state, particularly in the resting zone in the tibia (Figure 1B) and the bi-directional growth plate in the basal skull, the spheno-occipital synchondrosis (Figure 1C). EdU labeling revealed a significant increase in cell proliferation in resting zone chondrocytes but not in proliferating chondrocytes (Figure 1D).

**Figure 1.**
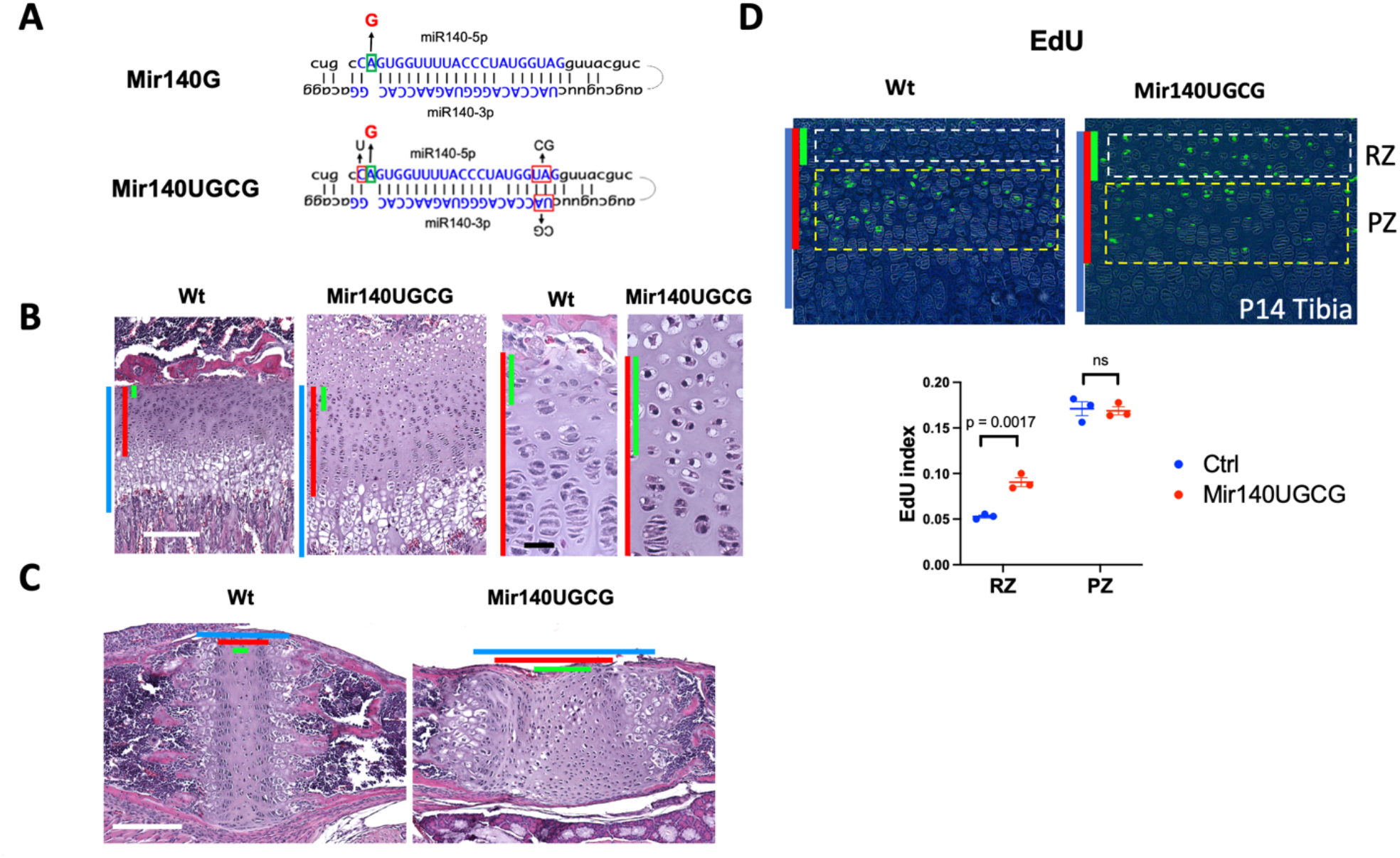
A gain-of-function mutation of miR-140-5p with enhanced expression causes expansion of resting zone chondrocytes in mice. (A) Generation of mice expressing a gain-of-function (GOF) mutant miR-140-5p (miR-140G) with enhanced expression. The indicated nucleotide changes were sequentially introduced into the mouse genome via i-GONAD. The GOF mutation (A to G) along with mutations at 5’ ends of -5p and -3p strands to increase the -5p abundance are indicated. (B) Hematoxylin and Eosin (H/E)-stained sections of the proximal tibial growth plate of P14 mice. The length of the growth plate (blue bars), the proliferating zone and resting zone (red bars), and the resting zone (green bars) is significantly expanded in heterozygous mutant mice (Mir140UGCG, *Mir140*^*UGCG/+*^). (C) H/E-stained sections of the spheno-occipital growth plate of the basal skull. The expansion of the resting zone is observed In this bidirectional growth plate. (D) EdU labeling of the tibial growth plate. Cellular proliferation is increased in the resting zone (RZ) in mutant mice but not in the proliferating zone (PZ).

### Decreased Glycolysis and Increased Oxidative Phosphorylation in Mir140UGCG Chondrocytes

To understand the mechanism by which the GOF mutant miR-140-5p (miR-140G) stimulates resting zone chondrocyte proliferation, we performed RNA-seq using primary rib chondrocytes of P7 mice of the previous mouse model without 5’ nucleotide changes ^3^. We found that the expression of genes whose products of which function is related to energy metabolism was significantly altered; those that regulate glycolysis was reduced, whereas those that regulate mitochondrial function and metabolism were reciprocally upregulated in a dose-dependent manner (Figure 2A, B). The upregulation of genes whose products are associated with glutaminolysis and lipolysis also suggested an increase in TCA cycle metabolism. Using primary chondrocytes, we performed the Seahorse analysis to assess the glycolysis and mitochondrial activity. Oxidative phosphorylation (OXPHOS) was assessed by real-time measurement of oxygen consumption rate (OCR) and glycolysis was assessed by extracellular acidification rate, a surrogate index for lactate production. The results supported the notion of reciprocal changes in glycolysis and mitochondrial metabolism, suggested by the RNA-seq data. (Figure 2C). In our previous study, we identified a few direct targets of miR-140G including *Hif1a*. Hif1a is a major regulator of cellular energy metabolism in hypoxic conditions by stimulating glycolysis and suppressing mitochondrial metabolism ^6; 12^. Thus, suppression of *de novo* targets of miR-140G directly and indirectly reduces glycolysis, a major mechanism for energy production in chondrocytes ^10^, and increase mitochondrial metabolism to compensate for the energy deficit.

**Figure 2.**
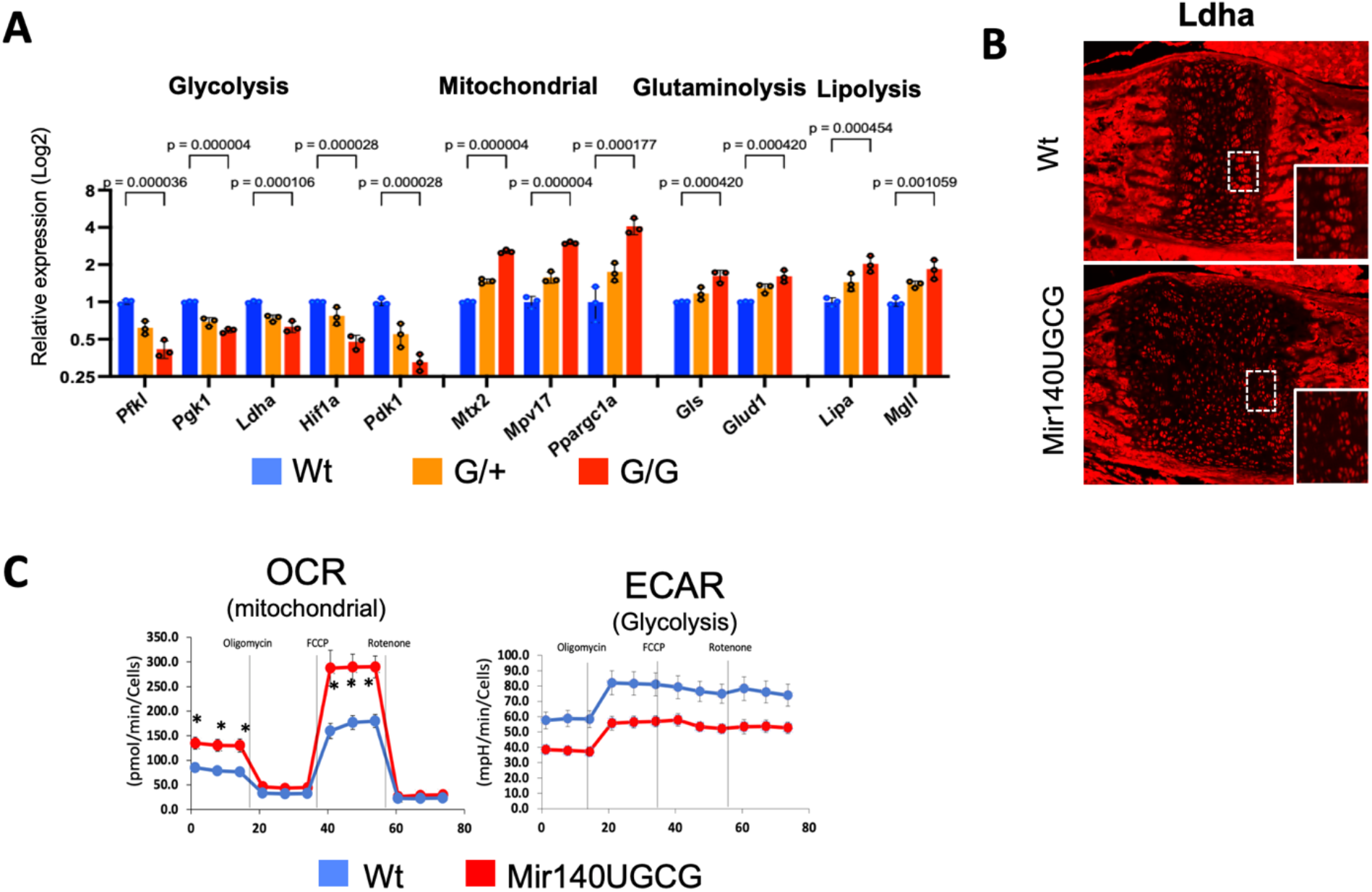
A gain-of-function mutation of miR-140-5p increase mitochondrial metabolism and decreases glycolysis in chondrocytes. (A) The expression of energy metabolism-related genes in primary rib chondrocytes from wildtype (Wt, *Mir140*^*+/+*^), heterozygous (G/+, *Mir140*^*G/+*^), and homozygous (G/G, *Mir140*^*G/G*^) mutant littermates. Gene expression data is obtained from the RNA-seq data ^3^. The expression of genes of which product function is related to glycolysis is down-regulated in a dose-dependent manner, whereas those related to mitochondrial metabolism were upregulated. +/- SEM, n = 3. (B) Downregulation of the Ldha, a key enzyme in the glycolysis pathway, is confirmed by immunofluorescence in the growth plate of the basal skull of Mir140UGCG (*Mir140*^*UGCG/+*^) mice. (C) Seahorse analysis of primary rib chondrocytes from P10 Mir140UGCG mice. Mutant chondrocytes show significant upregulation in oxygen consumption rate (OCR) with reciprocal downregulation of extracellular acidification rate (ECAR) suggesting an increase in mitochondrial metabolism and a reciprocal decrease in glycolysis. ±SEM, n = 8. * p < 0.001 in OCR, p < 0.001 at all points in ECAR, *t*-test

### Genetic Ablation of LDH Genes Increases Resting Chondrocyte Proliferation

Alterations of energy metabolism can have significant impacts on the regulation of cellular functions. Particularly, TCA cycle metabolites, such as acetyl CoA (Ac-CoA) and α - ketoglutarate (α KG), are known to play diverse regulatory roles. Thus, we hypothesized that altered energy metabolism contributed to the growth plate phenotype of *Mir140G* mutant mice. To test this hypothesis, we suppressed glycolysis by genetically ablating lactate dehydrogenase (LDH) genes. By catalyzing the conversion from pyruvate and lactate, LDH oxidizes nicotinamide adenine dinucleotide (NAD) + hydrogen (NADH) to generate NAD^+^, which is a co-substrate of Glyceraldehyde 3-phosphate dehydrogenase (GAPDH), one of the key enzymes of glycolysis. Therefore, reducing LDH action is expected to suppress glycolysis. To do so, we ablated *Ldha*, the most abundantly expressed LDH gene, in chondrocytes using *Col2-Cre* transgenic mice. A majority of *Col2-Cre:Ldha*^*fl/fl*^ (*Ldha* cKO) mice in the C57BL/6 background died at birth. However, in a congenic strain derived from the C57BL/6 and CD-1 backgrounds, *Ldha cKO* mice survive postnatally and grew into adulthood. *Ldha* cKO mice develop deformed and short long bones (Figure 3A). Histological analysis revealed bowing of the tibia, increased cellular density of the resting zone, and loss of flattening of proliferating columnar chondrocytes (Figure 3B). The proliferation of resting chondrocytes, but not proliferating columnar chondrocytes, was significantly increased in the *Ldha* cKO tibia (Figure 3C, D). In the spheno-occipital synchondrosis, a bidirectional growth plate in the basal skull, showed an expansion of the resting zone (Figure 3E). The expansion of the resting zone is also shared by *Mir140G* mutant growth plate, suggesting that the change in cellular metabolism is an underlying mechanism for the increased proliferation and expansion of resting chondrocytes.

**Figure 3.**
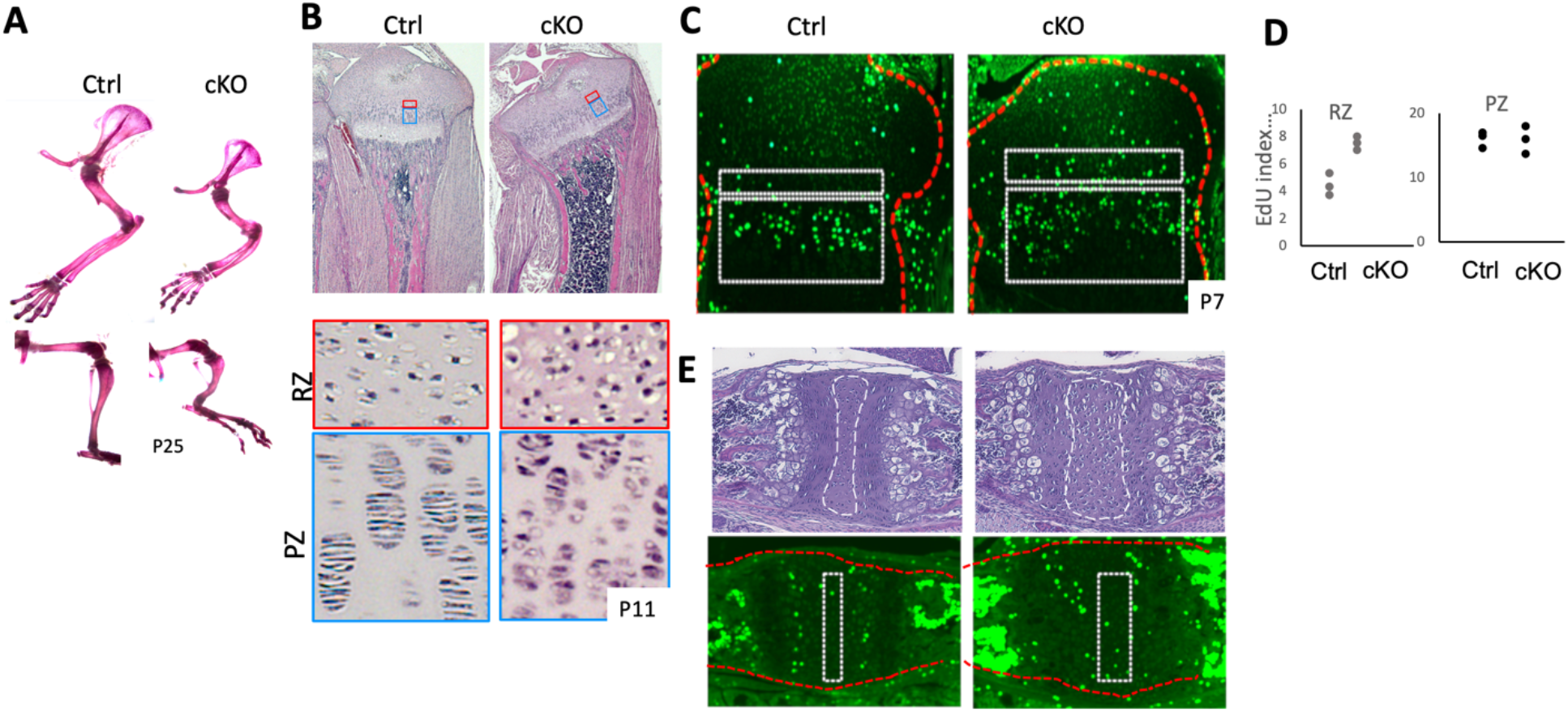
*Ldha*-deficiency in chondrocytes causes skeletal dysplasia. (A) Conditional deletion of *Ldha* in chondrocytes (cKO, *Col2-Cre:Ldha*^*fl/fl*^) causes shortening and bowing of limbs. Whole-mount alizarin red staining of fore- and hind limbs at 25 days of age. (B) Hematoxylin-Eosin (H/E)-stained sections of proximal tibial growth plates of P11 mice. The cellular density of the resting zone (RZ, indicated by red boxes) is increased in cKO mice. Cells in the columnar proliferating zone (PZ) show reduced flattening in cKO mice. (C - D) EdU staining in P7 tibial growth plates shows an increase in EdU-positive cells in the resting zone (top boxes in *C*, RZ in *D*) but not in the proliferating zone (bottom boxes in *C*, PZ in *D*). (E) H/E and EdU staining of the spheno-occipital growth plate of the basal skull. The resting zone is indicated by dotted boxes.

### Overlapping Role of Ldha and Ldhb

There are 4 genes that encode LDH in mice. According to our RNA-seq data ^3^, *Ldhb* is the second most abundantly expressed gene in chondrocytes next to *Ldha*, whereas *Ldhc* and *Ldhd* are hardly expressed. According to the International Mouse Phenotyping Consortium (IMPC, www.mousephenotype.org) database, *Ldhb*-null mice do not show overt skeletal abnormalities. To test whether *Ldhb* has an overlapping role with *Ldha* in chondrocytes, we generated *Ldhb*-null mice and cross them with *Ldha* cKO mice. An *Ldhb* germline-null mouse line was generated by i-GONAD in the congenic background as *Ldha* cKO mice. Both heterozygotes and homozygotes for *Ldhb-*null mice appear normal and fertile. However, compound mutant mice missing *Ldha* in chondrocytes and one allele of *Ldhb* (*Col2-Cre:Ldha*^*fl/fl*^*:Ldhb*^*+/-*^, *Ldh* compound *cKO*) showed a significantly smaller body size and growth defect compared with *Ldha* single conditional null mice (*Col2-Cre:Ldha*^*fl/fl*^), and they usually died by 3 weeks of age (Figure 4A, B). We never recovered live doubly homozygous null mice (*Col2-Cre:Ldha*^*fl/fl*^*:Ldhb*^*-/-*^), suggesting that complete loss of *Ldha* and *Ldhb* in chondrocytes results in embryonic or perinatal lethality. The shortening and bowing of long bones are more pronounced in *Ldh* compound *cKO* mice than *Ldha* single *cKO* mice (Figure 4C). Histologically, both *Ldha* cKO mice and compound mutant mice showed shortening of the growth plate, particularly the hypertrophic zone, and this phenotype is a little more severe in the compound mutants (Figure 4D). Using the *Ldh* compound *cKO* mutant mice, we performed Seahorse analysis to assess the impact of reduced LDH function in chondrocytes on energy metabolism. *Ldh cKO* chondrocytes showed mild downregulation of extracellular acidification rate (ECAR), a surrogate for lactate production, and reciprocal upregulation of oxygen consumption rate (OCR), an index of mitochondrial respiration.

**Figure 4.**
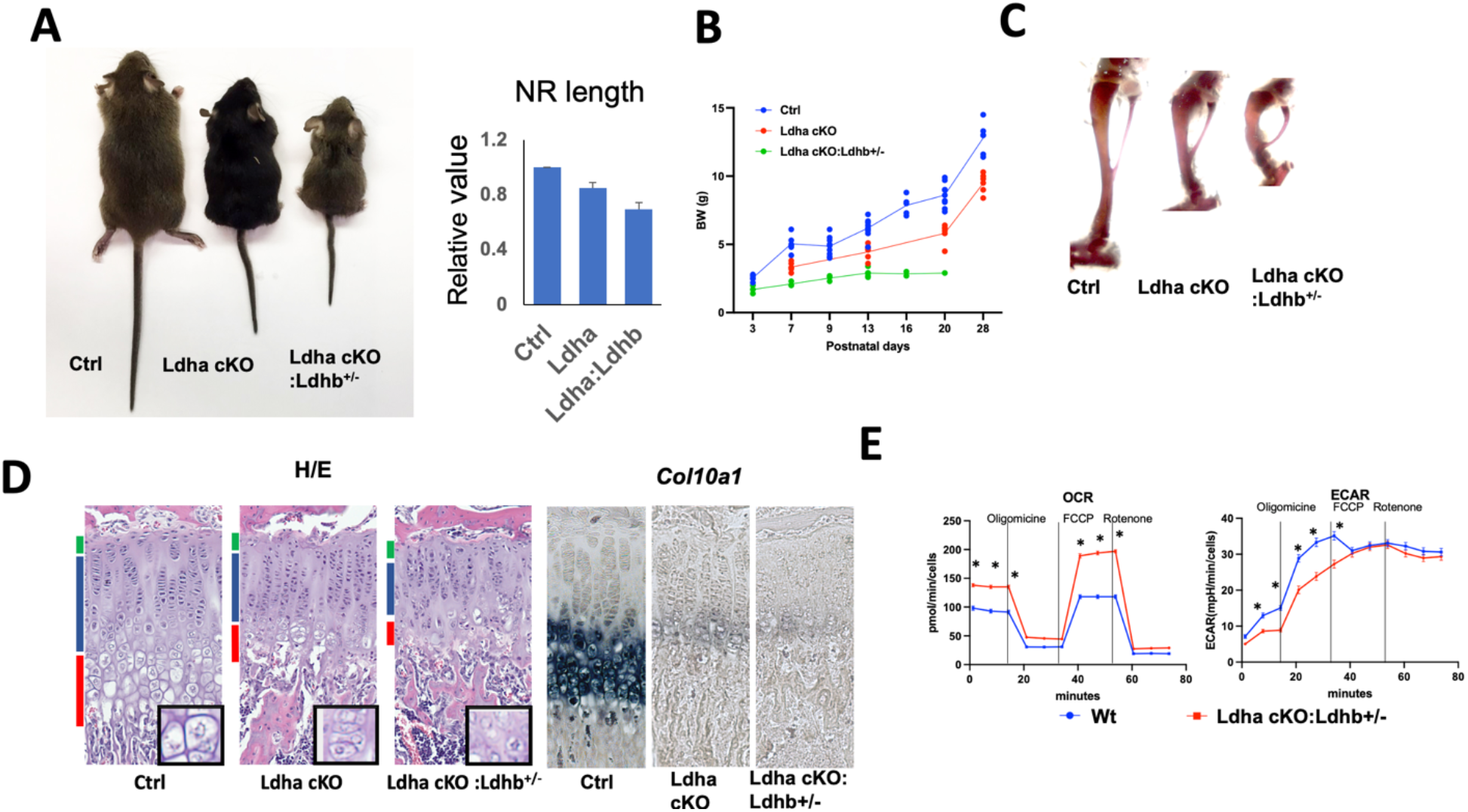
*Ldha* and *Ldhb* have overlapping roles in chondrocytes. (A - C) Additional heterozygous deletion of *Ldhb* causes significant growth defects in mice. Littermates of indicated genotypes at P21. *A* Gross appearance. *B* Nasal-ramp length at P14. *C* Whole-mount alizarin red staining of tibiae. *D* Weight gain curve. (D) H/E-staining and *Col10a1* expression of the proximal tibial growth plate. Insets show magnified views of the hypertrophic zone. The reductions in the number and in size of hypertrophic chondrocytes in *Ldha* cKO and *Ldha cKO:Ldhab*^*+/-*^ mice. The expression of *Col10a1* is also diminished in mutant mice. (E) Seahorse analysis of primary chondrocytes isolated from 10-day-old mice. A mild decrease in glycolysis and an increase in oxidative phosphorylation are suggested in mutant rib chondrocytes. ± SEM, n = 9, * p < 0.001, *t*-test.

### Reduction in Acetylation in Mir140UGCG and Ldh cKO Mice

The reduced glycolysis, a predominant mechanism that synthesizes ATPs in chondrocytes, likely led to compensatory upregulation of mitochondrial OXPHOS in *Mir140UGCG*, which then would increase the retention of TCA cycle metabolites in mitochondria and therefore reduces their export into the cytoplasm. Consistent with this notion, the cytoplasmic concentration of Acetyl-CoA (Ac-CoA), generated mainly from citrate derived from mitochondria, as well as the cytoplasmic citrate concentration, was significantly reduced in *Mir140UGCG* chondrocytes (Figure 5A). Because the availability of Ac-CoA in the cytoplasm and nucleus greatly influences acetylation of proteins, we evaluated the acetylation of histone proteins, the most abundant acetylated proteins in the cell (Figure 5B, C). As expected, the levels of acetylated histones in *Mir140UGCG* primary chondrocytes were reduced in mutant chondrocytes. Since a similar bioenergetic alteration is induced by *Ldha* ablation, we assessed histone acetylation in *Ldha* cKO chondrocytes. We also found reductions in histone acetylation in *Ldha-*deficient cells (Figure 5D, E).

**Figure 5.**
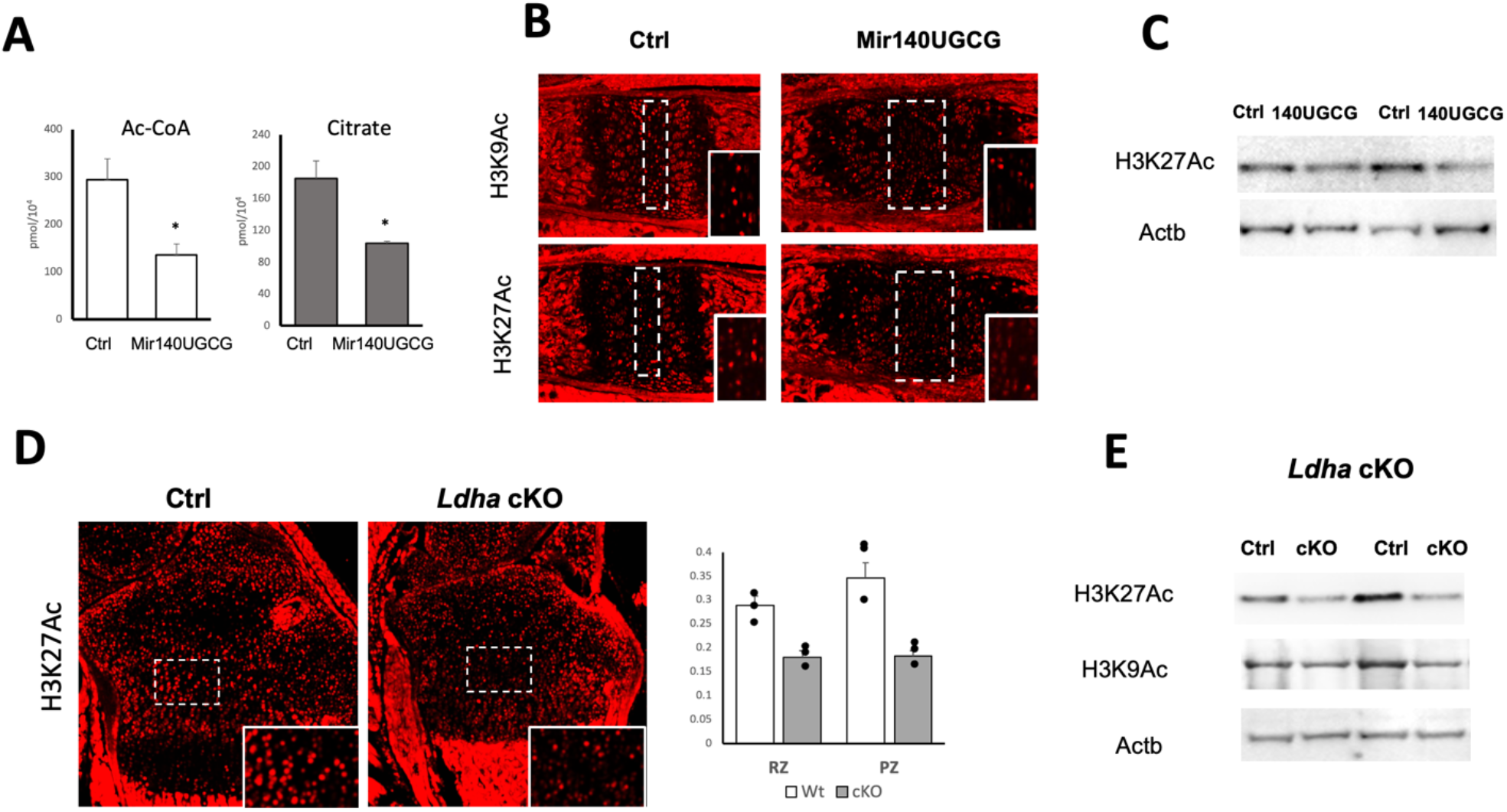
Altered energy metabolism affects histone acetylation. (A) Reduced cytosolic citrate and acetyl-CoA (Ac-CoA) levels in Mir140UGCG mice. Ac-CoA and citrate levels were determined in primary rib chondrocytes from 10-day-old mice. (B) Immunofluorescence of indicated acetylated histones in the spheno-occipital growth plate. Dotted boxes indicate the resting zone. Insets are magnified views of the resting zone. MIr1430UGCG mice show reduced histone acetylation. (C) Immunoblot analysis for H3K27 acetylation. (D) Immunofluorescence assessment of H3K27 acetylation in the tibial growth plate of control and *Ldha* cKO mice. Insets are magnified views of dotted boxes. (E) Quantification of H3K27 acetylation (H3K27Ac) in *D*. H3K27Ac-positive cells were counted in the resting zone and proliferating zone and normalized to the total number of cells per area. *Ldha* cKO show reduced numbers of chondrocytes positive for H3K27Ac. (F) Immunoblot quantification of acetylation of indicated histones in primary rib chondrocytes. *Ldha* cKO shows reduced histone acetylation.

### Skeletal Defects in Acly-Deficiency

Since the suppressed glycolysis is linked to reduced histone acetylation in *Mir140UGCG* and *Ldha* cKO mice, we hypothesized that reduced Acetyl-CoA (Ac-CoA) availability could be responsible for part of the skeletal phenotype in these mice. Cytoplasmic and/or nuclear Ac-CoA is generated from multiple sources, including citrate via the ATP-dependent citrate lyase, Acly, from dietary acetate via Acetyl-CoA synthases, and possibly from pyruvate via cytoplasmic/nuclear Pyruvate dehydrogenase (PDH), (PMID: 26039447). To mimic the citrate deficiency in *Mir140UCGC* mice, we conditionally ablated *Acly* in chondrocytes to reduce Ac-CoA availability. Conditional Acly-deficient (*Col2-Cre:Acl*^*fl/fl*^, *Acly* cKO) mice die at birth. The mutant long bones are short and bowed, which is reminiscent of the bone phenotype of *Ldha* or *Ldha:Ldhb* cKO mice (Figure 6A). Histological examination of the tibial growth plate also revealed the impairment of flattening of *Acly* cKO columnar proliferating as in *Ldha* cKO mice. In the spheno-occipital synchondrosis, *Acly* cKO mice show an expansion of resting chondrocytes, a growth plate phenotype shared by *Ldha* cKO and *Mir140UGCG* mice (Figure 6B). As expected *Acly-*deficient primary rib chondrocytes showed reduced levels of acetylated histones (Figure 6C). Interestingly, phosphorylation of AMPK was not altered in *Acly* cKO mice, unlike *Ldh* cKO chondrocytes, suggesting that ATP synthesis is not affected by *Acly*-deficiency (Figure 6D).

**Figure 6.**
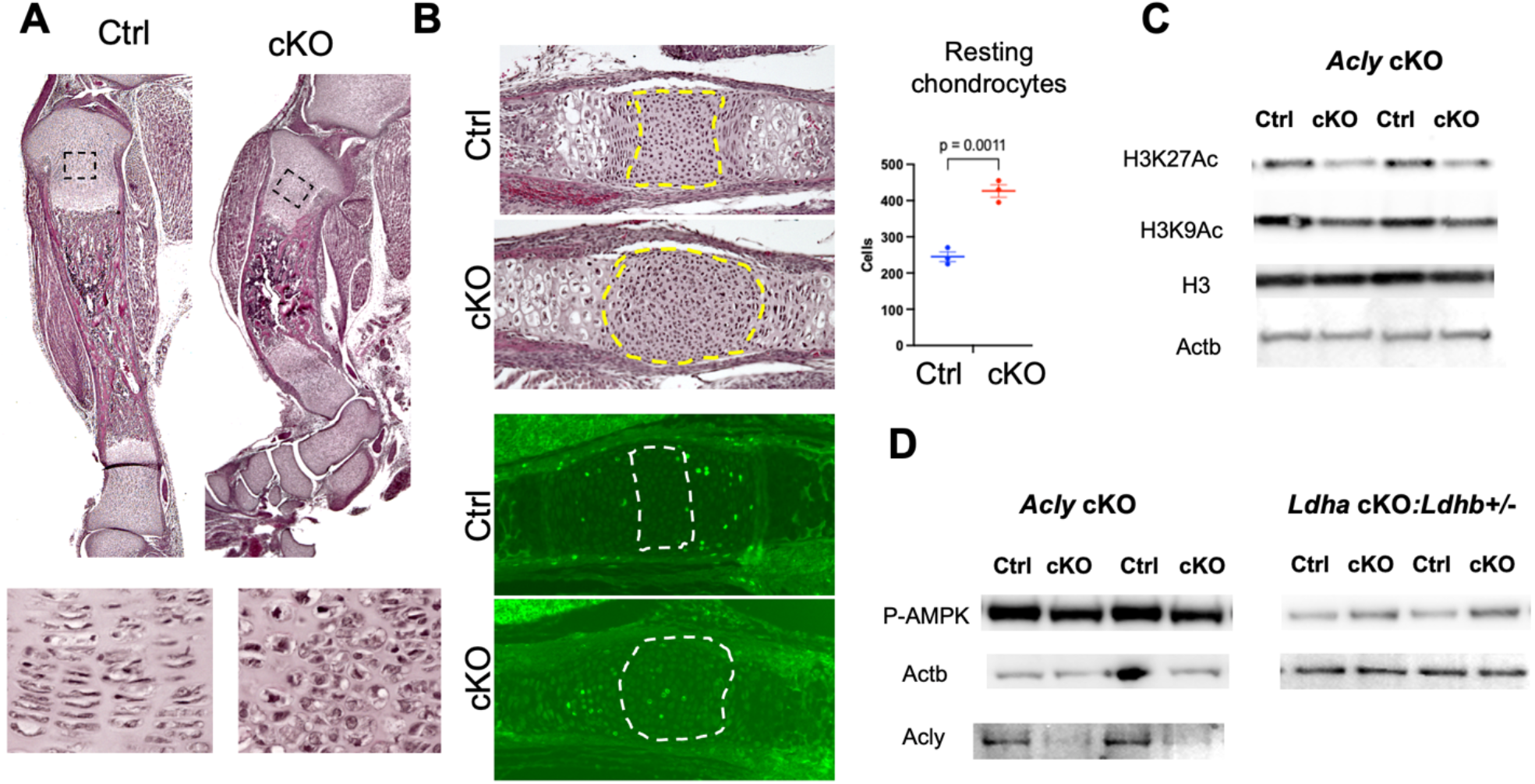
Skeletal defects of mice with conditional *Acly* deletion in chondrocytes. (A) *Acly* cKO mice show shortening and bowing of the tibia (*Top)*. Mice are perinatally lethal. H/E-stained sections of the tibia at P 0.5. The bottom panels show magnified views of proliferating chondrocytes in the boxed areas. The flat morphology of the columnar chondrocytes is lost in cKO mice. (B) Expansion of the resting zone chondrocytes in the spheno-occipital growth plate in E18.5 *Acly* cKO mice. *Left*, H/E-stained (*Top*) and EdU-stained (*Bottom*) sections. Boxed areas indicate the resting zone. *Right*, The number of resting chondrocytes is increased in the *Acly* cKO growth plate. (C) Reduced histone 3 acetylation in primary rib chondrocytes from E18.5 *Acly* cKO mouse embryos. Whole-cell lysates from an overnight culture of primary rib chondrocytes were subjected to immunoblot analysis. *Acly* cKO chondrocytes show reduced acetylation of H3K27 and H3K9. (D). Differential phosphorylation of AMPK between *Acly* cKO and *Ldh* compound cKO chondrocytes. The level of p-AMPK is unchanged in *Acly* cKO, whereas it is upregulated in *Ldh* compound cKO cells, suggesting ATP deficiency in *Ldh* compound cKO, but not in *Acly* cKO chondrocytes.

### Gene Expression Analysis in Ldh cKO and Acly cKO Chondrocytes

The phenotypic similarity between *Ldha* cKO and *Acly* cKO mice, including the shortening and bowing of long bones, expansion of resting chondrocytes, and reduced flattening of proliferating chondrocytes, prompted us to compare these two models at the molecular level. We performed RNA-seq analysis using P10-old primary rib chondrocytes of *Ldh* cKO chondrocytes and E18.5 *Acly* cKO rib chondrocytes. Because of the perinatal lethality of *Acly* cKO mice, we chose the late embryonic stage for chondrocyte isolation. Gene expression changes were evaluated by comparing mutants and littermate controls in these models. The average fold change of each gene was calculated in two models and plotted (Figure 7A). We found a mild direct relationship in the changes in gene expression between *Ldh* cKO and *Acly* cKO chondrocytes. Pathway analysis on the most significantly altered gene sets showed overlapping ontology terms between these two models, suggesting, *Ldh* and *Acly* gene perturbations cause similar molecular consequences (Figure 7B). Among these genes that are most significantly changed in both models, we found upregulations of groups of genes encoding chondrocyte-associated matrix proteins and downregulation of genes associated with endoplasmic reticulum (ER) stress (Figure 6C). Additionally, we found upregulation of Fgfr3 that was included in the group with the term, “PI3K/AKT signaling pathway”. Interestingly, the downregulation of ER stress genes and upregulation of Fgfr3 were also found in *Mir140G* chondrocytes.

**Figure 7.**
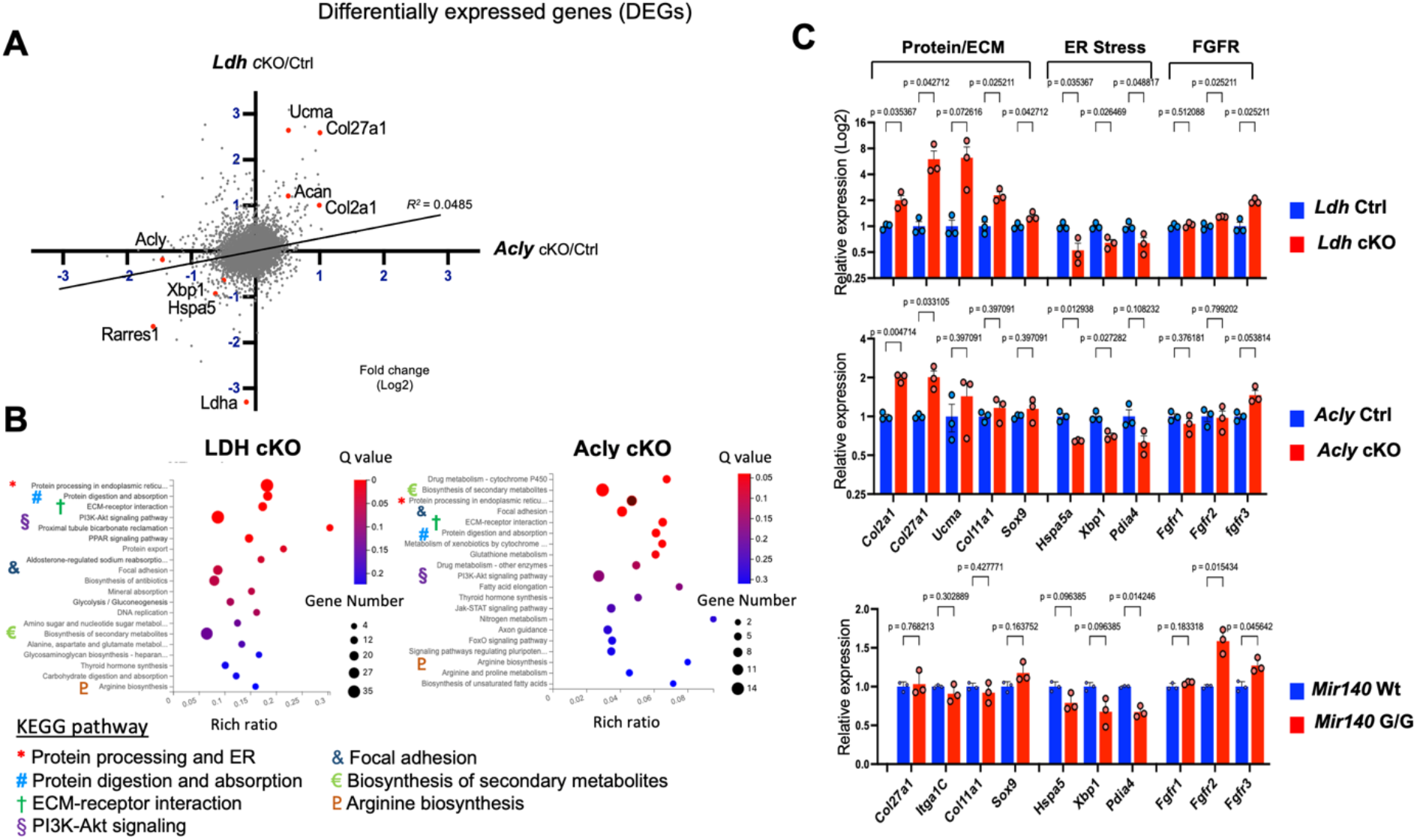
Gene expression analysis of *Ldh* cKO and *Acly* cKO chondrocytes. (A) Differentially expressed genes in *Ldh* cKO and *Acly* cKO chondrocytes. The fold change of genes in comparison between cKO and littermate control in two models, *Ldh* (Y axis) and *Acly* (X axis) is plotted. RNA, isolated from an overnight culture of primary rib chondrocytes of P5 *Col2Cre:Ldha*^*fl/fl*^*:Ldhb*^*+/-*^ (*Ldh* cKO) and littermate control mice, and from E18.5 *Col2-Cre:Acly*^*fl/fl*^ (*Acly* cKO) mice and littermate control were subjected to RNA-seq analysis. The gene expression changes caused by LDH and Acly ablation show a mild direct correlation. Simple linear regression model, R^2^ = 0.0485, p < 0.0001. (B) Pathway analysis on differentially expressed genes in *Ldh* cKO and *Acly* cKO chondrocytes show overlapping pathway terms indicated by colored symbols. The top 20 most significantly enriched terms are shown. (C) Differentially expressed genes associated with indicated terms. Genes encoding chondrocyte-specific matrix proteins are generally upregulated both in *Ldh* and *Acly* cKO chondrocytes, whereas those associated with the term “ER stress” are downregulated. Expression of *Fgfr3*, associated with the term, “PI3K/Akt signaling” is increased in *Ldh* and *Acly* cKO chondrocytes. Downregulation of “ER stress”-associated genes and upregulation of *Fgfr3* is also found in *Mir140G/G* chondrocytes. The gene expression data of *Ldh* and *Acly* cKO, and Mir140G/G is found at the Gene Expression Omnibus with Accession # GSE192971 and # GSE98309, respectively.

### Activation of Fgfr3 Signaling Causes Expansion of the Resting Zone

Activation mutations of FGFR3 cause achondroplasia, short-limb dwarfism. It is well known that activation of FGF signaling decreases the proliferation of proliferating chondrocytes. However, it was previously suggested that FGF signaling stimulates chondrocyte proliferation during embryonic stages. This could be explained by that FGF signaling could have opposite effects on resting chondrocytes and proliferating chondrocytes. To investigate the effect of FGF signaling on resting chondrocytes, we overexpressed a constitutively active mutant Fgfr3 (Fgfr3 K650E, Fgfr3*) in chondrocytes. Both embryonic Fgfr3* expression using a *Col2-Cre* driver and perinatal Fgfr3* expression at E18.5 using a tamoxifen-inducible chondrocyte-specific Cre driver (*Sox9-CreER*) showed significant expansions of the resting zone, whereas it reduces the size of proliferating chondrocytes, demonstrating that FGFR3 signaling has a different effect on resting chondrocytes from proliferating chondrocytes (Figure8 A-C).

**Figure 8.**
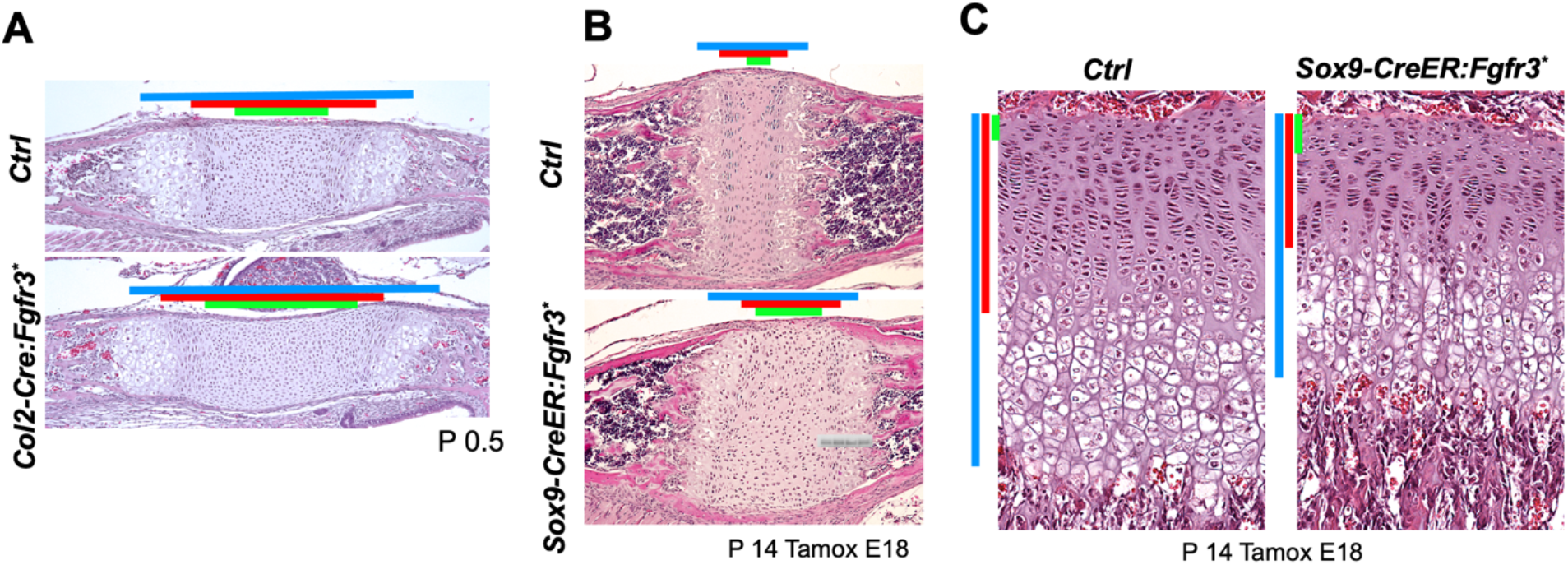
Differential effects of overexpression of a constitutively active Fgfr3 on resting and proliferating chondrocytes. (A - C) A constitutively active Fgfr3 (Fgfr3^K650E^) is overexpressed from the *Rosa26* locus (Chr. 6q) in chondrocytes using the constitutively active, chondrocyte-specific *Col2-Cre* driver (A) or the tamoxifen-inducible *Sox9-Cre-ER* driver (B, C). In both models, the resting zone of the spheno-occipital synchondrosis (A, B) is expanded upon overactivation of Fgfr3 signaling, whereas it reduces the proliferating zone as best appreciated in the tibia (C). Colored bars indicate, the total growth plate length (blue), the combined length of the resting and proliferating zones (red), and the resting zone (green).

### Effects on Lipid Metabolism

Ac-CoA deficiency can also negatively affect lipid synthesis. We extracted lipid species from primary *Acly* cKO and *Mir140G* chondrocytes and subjected them to mass spectrometry analysis (Figure 9A, B, Supple Table). Each lipid species was quantified and categorized into 6 groups, and the fold differences between control and mutant chondrocytes were plotted. Most lipid groups except the cholesterol ester group were not much different. Cholesterol esters, mostly present as lipid storage in the cell, were more abundant in *Acly* cKO chondrocytes and were less abundant in *Mir140UGCG* chondrocytes. This may be due to the possible increase in beta-oxidation in *Mir140UGCG* chondrocytes to compensate for the impaired ATP synthesis. Ras prenylation can be one of the indicators for the *de novo* lipid synthesis pathway, as prenyl pyrophosphates are intermediate metabolites in the cholesterol synthesis pathway, and protein prenylation plays an important regulatory role in several signaling pathways ^27^. Interestingly, Ras prenylation was unaffected in *Acly* or *Ldh* cKO chondrocytes while Lovastatin treatment that suppresses the cholesterol synthesis pathway reduced it in primary chondrocytes (Figure 9C). This result suggests that alternative pathways, particularly the Acss2-mediated Ac-CoA synthesis from acetate, compensate for the reduced Ac-CoA synthesis from citrate in these mutant chondrocytes for lipid synthesis reasonably well but less so for histone acetylation.

**Figure 9.**
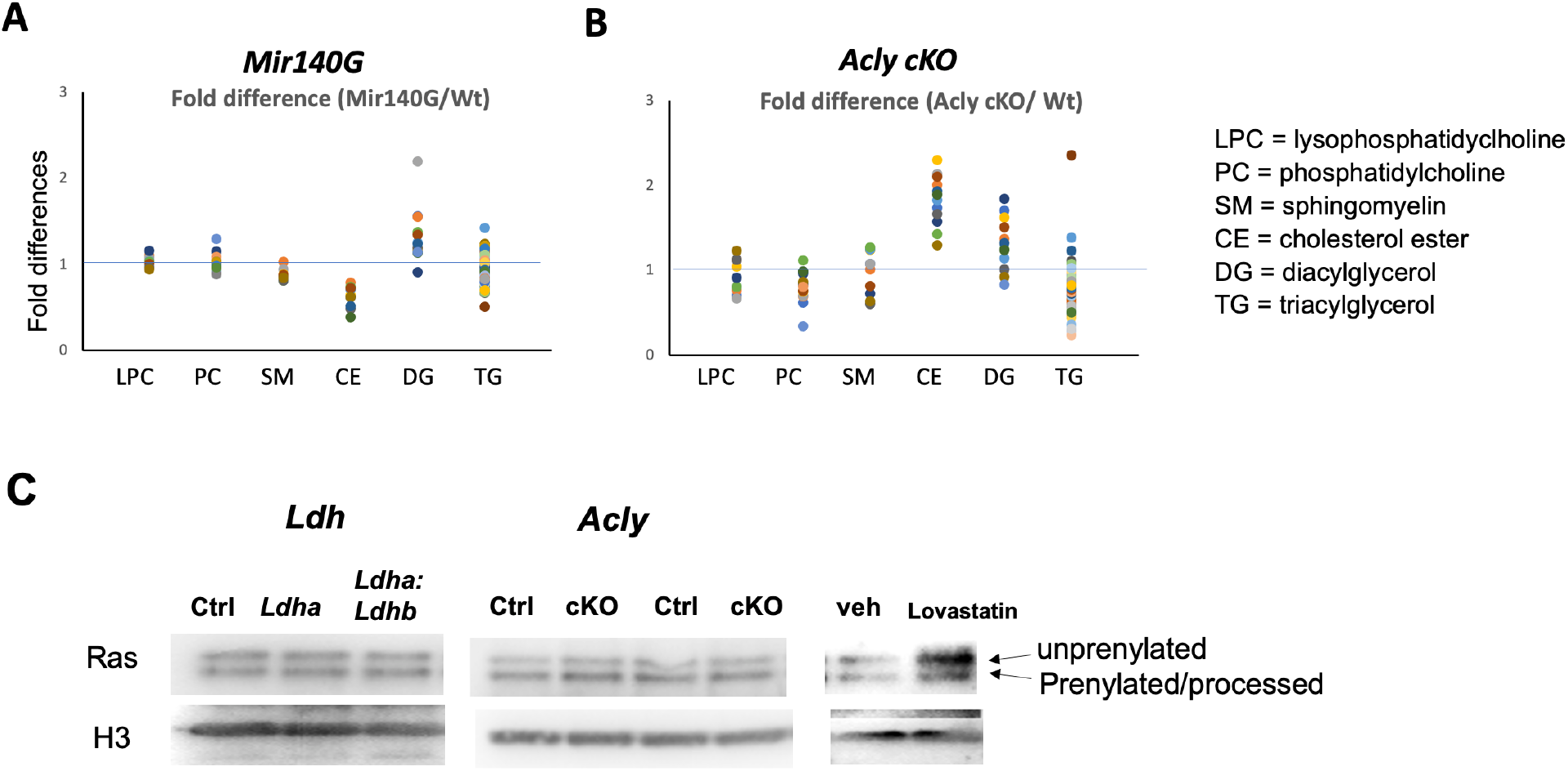
Lipid metabolism is not significantly altered in *Acly* cKO or *Mir140UGCG* chondrocytes. (A, B) Comparison of 6 groups of lipid species between control and mutant chondrocytes. Lipid quantification of culture cells was performed by mass spectrometry analysis. Values were normalized by the cell mass, and the fold difference of each lipid species is plotted. The abundance of most lipid species appears generally unchanged except that cholesterol esters are reduced in *Mir140UGCG* chondrocytes (A), whereas they are increased in *Acly* cKO chondrocytes (B). (C) Ras prenylation was assessed by immunoblot analysis using whole cell lysates. Prenylated Ras proteins migrate faster due to subsequent processing. Whereas Lovastatin treatment to suppress the cholesterol synthesis pathway decreased prenylation, it was unaffected in *Ldh-* or *Acly*-deficient chondrocytes.

## Discussion

In this study, we show that the disease-causing GOF mutant miR-140 decreases glycolysis and increases OXPHOS in growth plate chondrocytes, that reduced glycolysis via *Ldh* ablation increases resting chondrocyte proliferation and reduces histone acetylation, and that *Acly* disruption causes cellular and molecular abnormalities similar to those of *Ldh* ablation in chondrocytes. These findings suggest that suppression of glycolysis stimulates resting chondrocyte proliferation by reducing the cytoplasmic/nuclear Ac-CoA availability.

The mechanism by which the proliferation of resting chondrocytes is regulated is poorly understood. *Mir140G/G* and *Mir140UGCG* mouse models with the unique growth plate phenotype, *i*.*e*., expansion of the resting zone, provided an opportunity to investigate the regulatory mechanisms that control resting chondrocyte proliferation. It was shown that Indian hedgehog (Ihh) signaling increases the proliferation of resting chondrocytes ^28^. However, it does not cause an expansion of the resting zone; therefore the increase in proliferation may be a secondary consequence of the stimulated differentiation of resting chondrocytes into columnar proliferating chondrocytes ^29^. In non-chondrocytic skeletal progenitor cells, *Ras* signaling stimulates the proliferation of *Col2*-positive osteochondro-progenitors in the bone marrow ^30^. Our data that activation of Fgfr3 signaling causes an expansion of the resting zone of the growth plate supports the idea that the Ras signaling pathway might be a common regulator of skeletal stem/progenitor cell proliferation.

The findings from *Mir140G, Ldh* cKO, and *Acly* c KO mice, suggest a regulatory role of energy metabolism in controlling resting chondrocyte proliferation and differentiation of columnar proliferating chondrocytes. Because *Acly* cKO chondrocytes do not show increases in phospho-AMPK unlike *Mir140G* or *Ldh* cKO chondrocytes, energy deficiency is an unlikely cause for the phenotype. On the other hand, reduced histone acetylation was commonly observed in these three models, suggesting that the decreased Ac-CoA availability is responsible for the phenotype. Ac-CoA serves as the donor of the acetyl group for protein modification and also as a precursor molecule for *de novo* lipid synthesis. Our lipid quantification and assessment of Ras prenylation to indirectly assess the cholesterol synthesis pathway in primary chondrocytes from these mice showed no overt deficits, suggesting that lipid synthesis is not severely affected, unlike histone acetylation.

The increased expression of Fgfr3 commonly observed in these models and the stimulatory effect of Fgfr3 signaling in resting zone chondrocytes are consistent with the hypothesis that upregulation of Fgfr3 can be the underlying cause for the expanded resting zone. However, the mechanism by which Ac-CoA deficiency can lead to the *Fgfr3* upregulation and these phenotypic changes is not clear at the moment. The decrease in histone acetylation suggests that the reduced availability of Ac-CoA could epigenetically alter gene expression including *Fgfr3* or functions of undetermined regulatory proteins. For example, Hif1a acetylation regulates its stability and transcriptional activity ^31; 32^.

Together, these results suggest that chondrocyte bioenergetics regulate gene expression by changing Ac-CoA metabolism, which differentially controls resting and proliferating chondrocytes possibly through FGFR3 signaling (Figure 10A, B). Currently, firm evidence for upregulation in FGFR3 signaling or epigenetic deregulation of the Fgfr3 gene in these mouse models is missing; the proposed mechanism remains hypothetical.

**Figure 10.**
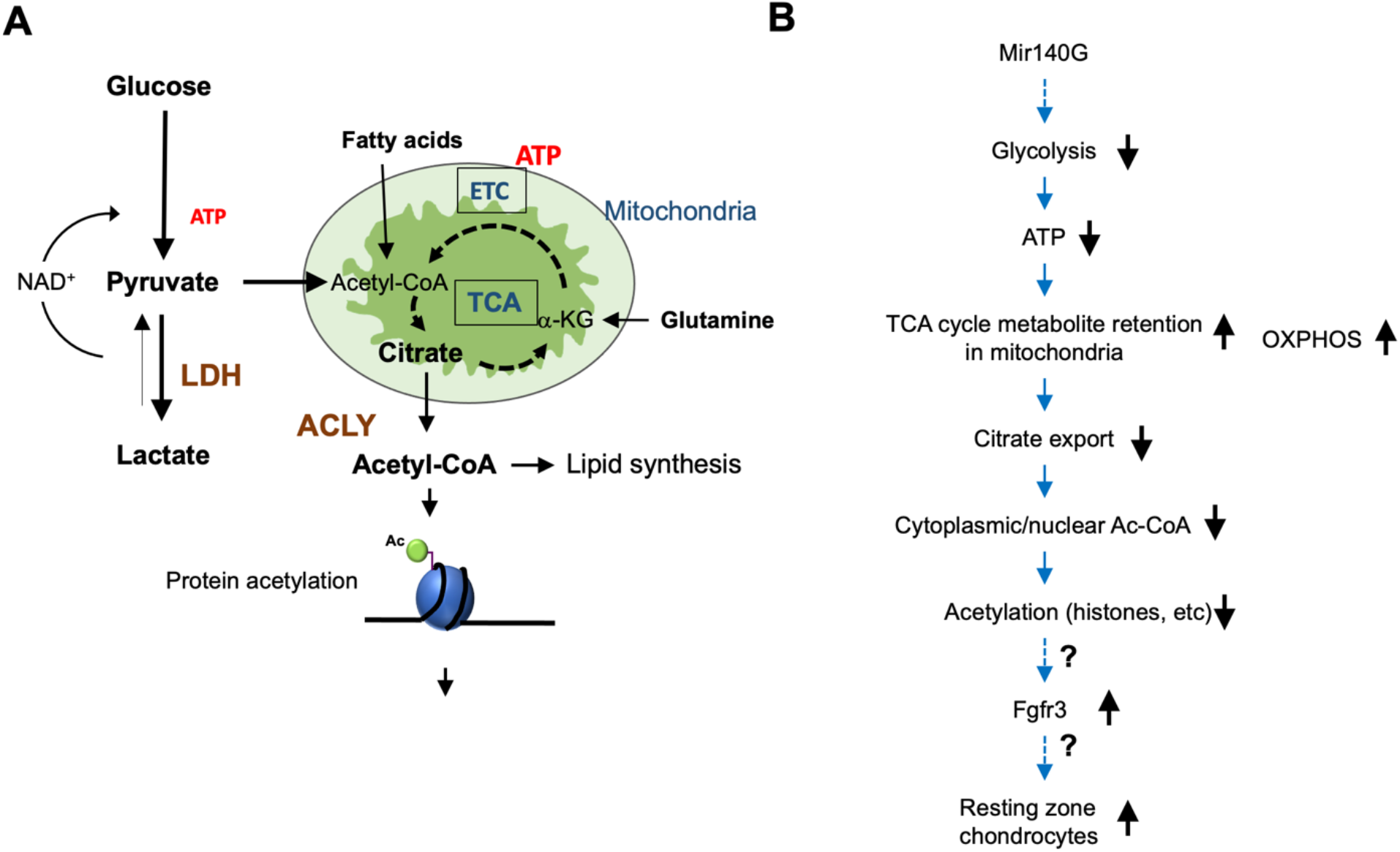
Hypothetical model linking energy metabolism and regulation of chondrocytes. (A) Schematic representation for energy metabolism pathways. (B) Hypothetical model in which *Mir140G* causes expansion of resting chondrocytes.

## Supporting information

Lipid analysis

## Data availability

GEO GSE192971

## Acknowledgements

We thank the center for skeletal research (P30, AR066261, AR07042) for assistance of histological analysis. This project was funded by the NIH grant AR056645. We have no conflicts of interest to declare.

